# Host EV–derived lipids promote malaria parasite development via mosquito fatty acid β-oxidation

**DOI:** 10.1101/2025.11.09.687433

**Authors:** Xiumei Song, Weimin Cai, Jiao Wu, Lijuan Liu, Xinran Gu, Han Zhou, Hui Zhao, Penghai Tian, Qian Xu, Kun Qian, Wei Huang, Jingwen Wang

## Abstract

Malaria parasites must transition between mammals and mosquitoes, but how they adapt to such distinct environments is unclear. We show that blood from infected mammals buffers this environmental shift by packaging parasite-induced signals into extracellular vesicles (EVs). EVs internalized by mosquito midgut increases expression of VLCAD and raises levels of a 22-carbon fatty acid, driving fatty acid breakdown and acetyl-CoA production. The parasite exploits this metabolic shift to synthesize its own lipids during oocyst growth. Stable isotope tracing confirmed that^13^C_4_-labelled behenic acid is degraded and incorporated into parasite fatty acids. Lipidomic profiling further revealed that ceramides enriched in the EVs are key mediators. These findings identify a mechanism by which host bood-derived EVs reprogram mosquito lipid metabolism to promote parasite transmission.

## Introduction

Malaria, caused by *Plasmodium* spp. and transmitted by *Anopheles* mosquitoes, remains a major global health challenge. Despite decades of control efforts, malaria cases resurged to 263 million in 2023, exceeding incidence levels of the past decade^1^. *Plasmodium* parasites alternate between mammalian and mosquito hosts, confronting dramatic shifts in temperature, oxygen, nutrient availability, and immune pressure. The mechanisms that enable these adaptations remain poorly understood.

A major bottleneck occurs during transmission to the mosquito midgut. Following a blood meal, *Plasmodium* gametocytes are ingested together with host blood and undergo rapid morphological transitions, from gametocytes to gametes, zygotes, ookinetes, and eventually early oocysts, within ∼48 hrs. During this time, blood digestion releases metabolites that fuel mosquito reproduction^2,3^ and simultaneously shape parasite development. For example, the blood-derived metabolite xanthurenic acid induces gamete exflagellation^4^, human insulin suppresses mosquito immune signaling to promote *Plasmodium falciparum* (*P. falciparum*) infection^5^, and reduced mammalian serotonin dampens mitochondrial ROS production to enhance parasite survival^6^. These findings suggest that the blood meal serves not only to nourish the mosquito but also to buffer parasites against the sudden metabolic and environmental shifts encountered in the midgut.

In addition to soluble metabolites, host serum contains EVs, which are emerging regulators of intercellular communication. In mammalian hosts, *P. falciparum* modifies EVs released from infected red blood cells (iRBCs), enabling parasite–parasite and parasite–host communication. For example, iRBC-derived EVs transfer DNA plasmids to promote drug resistance and gametocytogenesis^7^, activate NF-κB in splenic fibroblasts to upregulate ICAM-1^8^, impair endothelial barrier integrity via miRNA– Argonaute 2 complexes^9^, and remodel naïve red blood cells by delivering 20S proteasome complexes^10^. These activities collectively enhance parasite survival and invasion. However, whether mammalian blood-derived EVs modulate *Plasmodium* transmission within mosquitoes remains unknown.

Here, we show that EVs from *Plasmodium*-infected blood enhance mosquito infection by reprogramming lipid metabolism in the midgut. EVs uptake upregulates VLCAD, boosting β-oxidation of C22 fatty acids and generating acetyl-CoA that is subsequently exploited by the parasite for fatty acid synthesis and oocyst growth of both *P. berghei* and *P. falciparum*. Lipidomic profiling identified ceramides as key EVs cargo sufficient to promote parasite infection. Collectively, our findings uncover a previously unrecognized role for host blood-derived EVs in enhancing *Plasmodium* transmission by rewiring mosquito fatty acid metabolism.

## Results

### EVs from *Plasmodium*-infected blood promote oocyst development

To determine the influence of blood-derived EVs on parasite infection in mosquitoes, we isolated EVs from the sera of *P. berghei* infected (EV^infected^) and uninfected (EV^normal^) mice by ultracentrifugation and orally supplemented them to mosquitoes through sugar feeding (Fig. 1A). Nanoparticle tracking analysis (NTA) and transmission electron microscopy (TEM) revealed no significant differences in particle number or morphology between EV^infected^ and EV^normal^ (Fig. S1A and S1B). To test whether host-derived EVs are internalized by mosquito cells, we labelled EVs with PKH67, a lipophilic fluorescent dye commonly used for EVs tracking^11^. PKH67 signals were detected in the cytoplasm of MSQ43 mosquito cells *in vitro* (Fig. S2A) and in mosquito midgut *in vivo*, with comparable fluorescence intensities between EV^infected^ and EV^normal^ (Fig. 1B). No EV uptake was observed in ovaries or malpighian tubules (Fig. S2B).

**Fig. 1.**
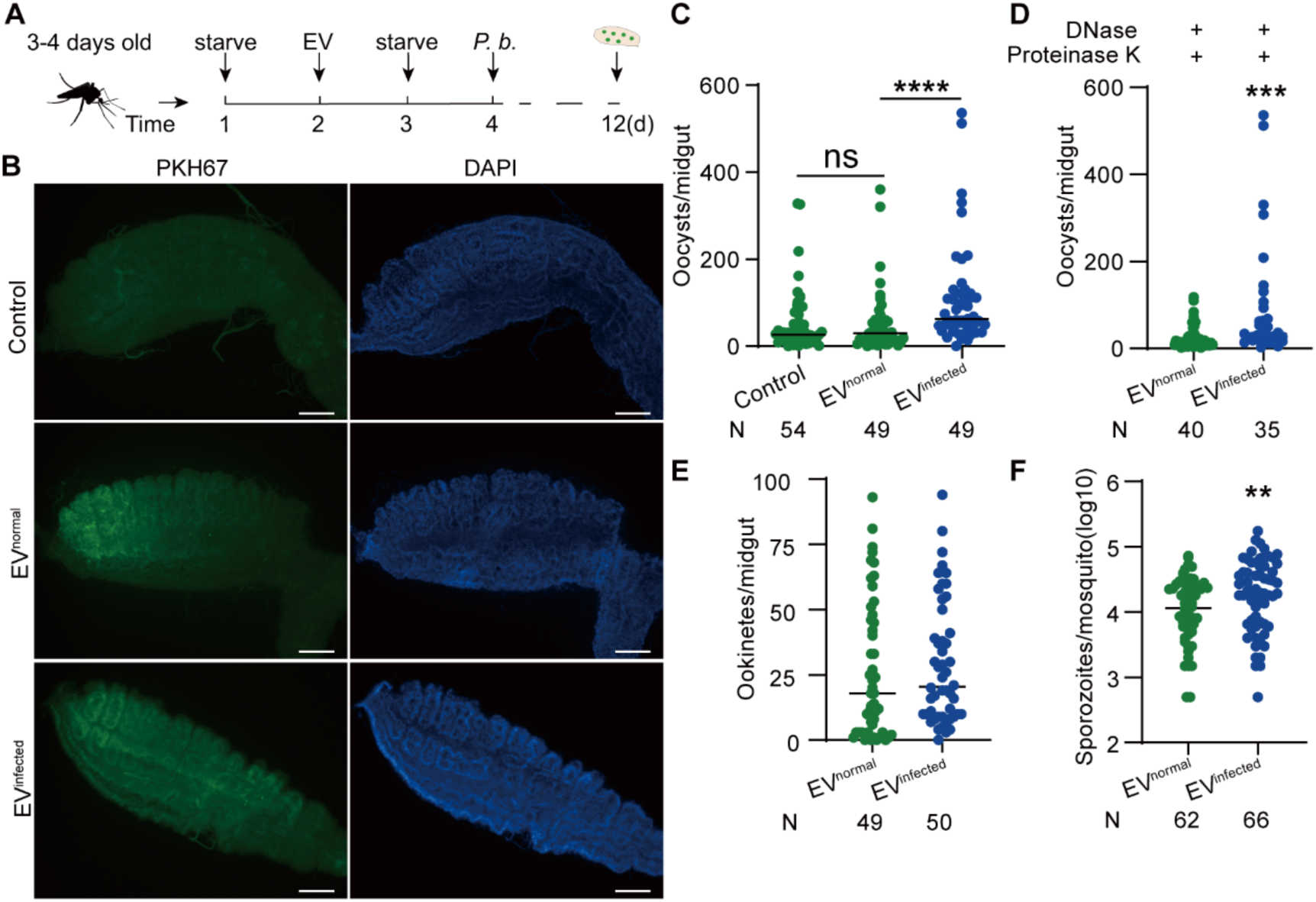
EVinfected promotes *P. berghei* infection in mosquitoes. **A**, The workflow of EVs treatments in *An. stephensi*. **B**, Internalization of EV^normal^ and EV^infected^ labelled with PKH67 (green) in mosquito midgut. Nucleis were stained by DAPI (blue). Scale bar, 50 μm. **C**, Oocyst numbers of mosquitoes treated with EV^normal^, EV^infected^ and non-EVs control. **D**, Oocyst numbers of mosquitoes orally supplemented with DNase and Proteinase K-treated EV^normal^ and EV^infected^. **E** and **F**, Ookinete **(E)** and sporozoite **(F)** numbers of mosquito treated with EV^normal^ and EV^infected^. For (C) to (F), data were pooled from two independent experiments, and each dot represents an individual mosquito. N, number of mosquitoes analyzed. The horizontal black lines indicate the median number of oocysts. Significance was determined by ANOVA with Dunn’s test in (C) and Mann-Whitney test in (D) to (F). \*\**p* < 0.01, \*\*\**p* < 0.001, \*\*\*\**p* < 0.0001. ns, not significant.

Functionally, oral administration of EV^infected^ significantly increased oocyst numbers compared to EV^normal^ or non-EV controls (Fig. 1C). To exclude potential effects of co-isolated contaminants, we treated the isolated EV^infected^ and EV^normal^ with proteinase K and RNase to remove protein and RNA contaminants and confirmed that EV^infected^ still promoted oocyst development (Fig. 1D). To identify the parasite stage influenced by EV^infected^, we quantified ookinetes, a stage prior to oocyst, and found no difference between treatment groups (Fig. 1E). We also examined the sporozoite numbers in the salivary glands and found that a significant increase in sporozoite numbers in EV^infected^-treated mosquitoes (Fig. 1F), indicating that EVs play a crucial role in promoting *Plasmodium* oocyst maturation in mosquitoes.

We next investigated whether EV^infected^ indirectly enhanced infection by altering mosquito feeding or reproduction. Neither blood meal size nor fecundity was affected by EVs supplementation (Fig. S3A and S3B). Altogether, these findings indicate that blood-derived EVs are specifically internalized by the mosquito midgut, where EV^infected^ enhance *P. berghei* oocyst development and transmission potential.

### EV^infected^ induces VLCAD expression to promote parasite development

To investigate how EV^infected^ promotes oocyst development, we performed proteomic profiling of mosquito midguts supplemented with EV^infected^ and EV^normal^ (Fig. 2A). Principal components analysis (PCA) analysis revealed clear separation between groups, indicating distinct protein expression signatures (Fig. 2B). In total, 63 proteins were differentially expressed, with 40 upregulated and 23 downregulated in EV^infected^-treated midguts (Fig. 2B). The strong induction of specific proteins suggested candidate mediators of parasite development. To test this, we individually knocked down the four genes upregulated more than sixflod: GTP: AMP phosphotransferase AK3 (GAK3), VLCAD, endoplasmic reticulum–Golgi intermediate compartment protein 3 (ERGIC3), and vacuolar protein sorting–associated protein 13D (VPS13D) (Fig. 2C). Notably, only *VLCAD* knockdown significantly reduced both oocyst and sporozoite numbers (Fig. 2D to 2F and S4). We next examined whether infectious blood itself induces VLCAD expression. Indeed, ingestion of *P. berghei*–infected blood (IB) significantly increased VLCAD transcript and protein levels in mosquito midguts compared with those from mosquitoes fed on non-infected mice (NB) (Fig. 2G and 2H). Together with the observation that EV^infected^ upregulates VLCAD, these data suggest that EVs are the active blood-derived factor driving VLCAD induction, thereby facilitating *P. berghei* infection in *An. stephensi*.

**Fig. 2.**
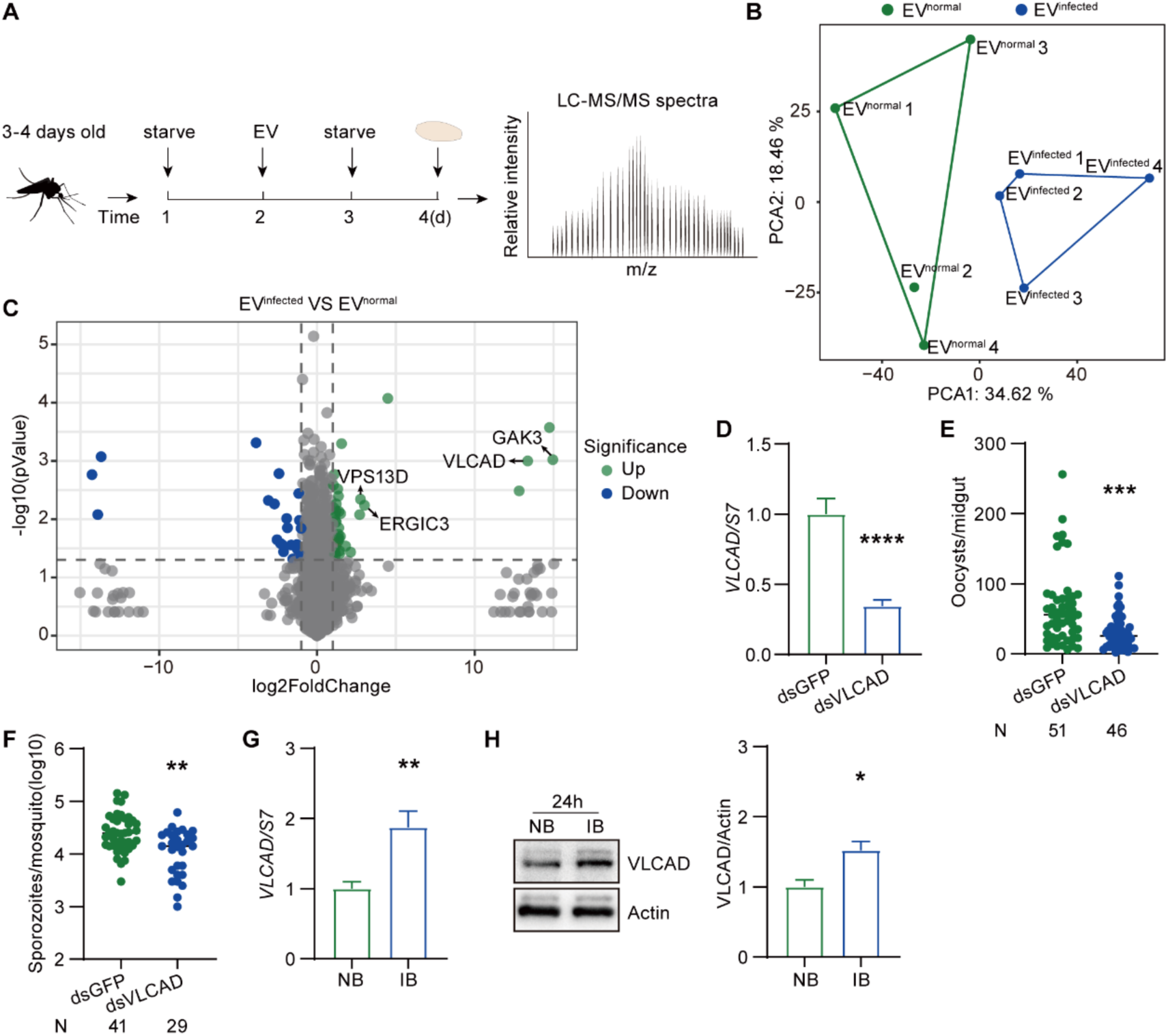
EVinfected induced-VLCAD promotes *Plasmodium* infection. **A**, The workflow of proteome analysis in the midgut of mosquitoes. **B**, PCA of proteins expression in the midgut of mosquitoes treated with EV^normal^ and EV^infected^. **C**, Volcano plot of the differentially expressed proteins between mosquitoes treated with EV^normal^ and EV^infected^. **D**, *VLCAD* knockdown efficiency in mosquitoes. The expression level of *VLCAD* was normalized to *An. stephensi S7*. Relative expression level of *VLCAD* in dsVLCAD mosquitoes was normalized to that in dsGFP controls. Data are presented as the mean ± SEM (n=10 in dsGFP, n=10 in dsVLCAD). **E and F**, Oocyst (**E**) and sporozoite (**F**) numbers in dsGFP and dsVLCAD –treated mosquitoes. Data were pooled from two independent experiments, and each dot represents an individual mosquito. N, number of mosquitoes analyzed. The horizontal black lines indicate the median number of oocysts. **G**, qPCR analysis of *VLCAD* levels in the midgut of mosquitoes fed on NB and IB. The expression level of *VLCAD* was normalized to *An. stephensi S7*. Relative expression level of *VLCAD* in the midgut of mosquitoes post IB were normalized to that of mosquitoes post NB. (n=10 in NB, n=10 in IB). Data are presented as the mean ± SEM. **H**, Western blot of VLCAD in the midgut of mosquitoes fed on NB and IB. The right panel was the quantification of band intensities of Western blot. The expression level of the target protein was normalized to Actin. Data were pooled from four independent experiments. Data are presented as the mean ± SEM. Significance was determined by two-sided Student’s *t* test in (D), (G) and (H) and Mann-Whitney test in (E) and (F). \**p* < 0.05, \*\**p* < 0.01, \*\*\**p* < 0.001, \*\*\*\**p* < 0.0001.

### VLCAD-dependent fatty acid catabolism drives parasite infection

VLCAD catalyzes the initial dehydrogenation of long-chain acyl-CoA (C14–C20) in mammalian mitochondria^12^. To investigate its function in *An. stephensi*, we profiled fatty acids in *VLCAD* knockdown mosquitoes two days after a blood meal using gas chromatography–mass spectrometry (GC–MS). Among 34 detected fatty acids, several very-long-chain fatty acids (VLCFAs), including C20:4n6, C22:0, C22:5n3, C22:5n6, accumulated significantly in dsVLCAD treated-mosquitoes compared to dsGFP treated-controls (Fig. 3A), indicating a conserved role for VLCAD in VLCFAs catabolism. We next asked whether the upregulation of VLCAD expression following EV^infected^ uptake was due to fatty acids accumulation. GC–MS analysis of midguts revealed elevated levels of C22:5n6 in EV^infected^– treated mosquitoes compared to EV^normal^ controls (Fig. 3B).

**Fig. 3.**
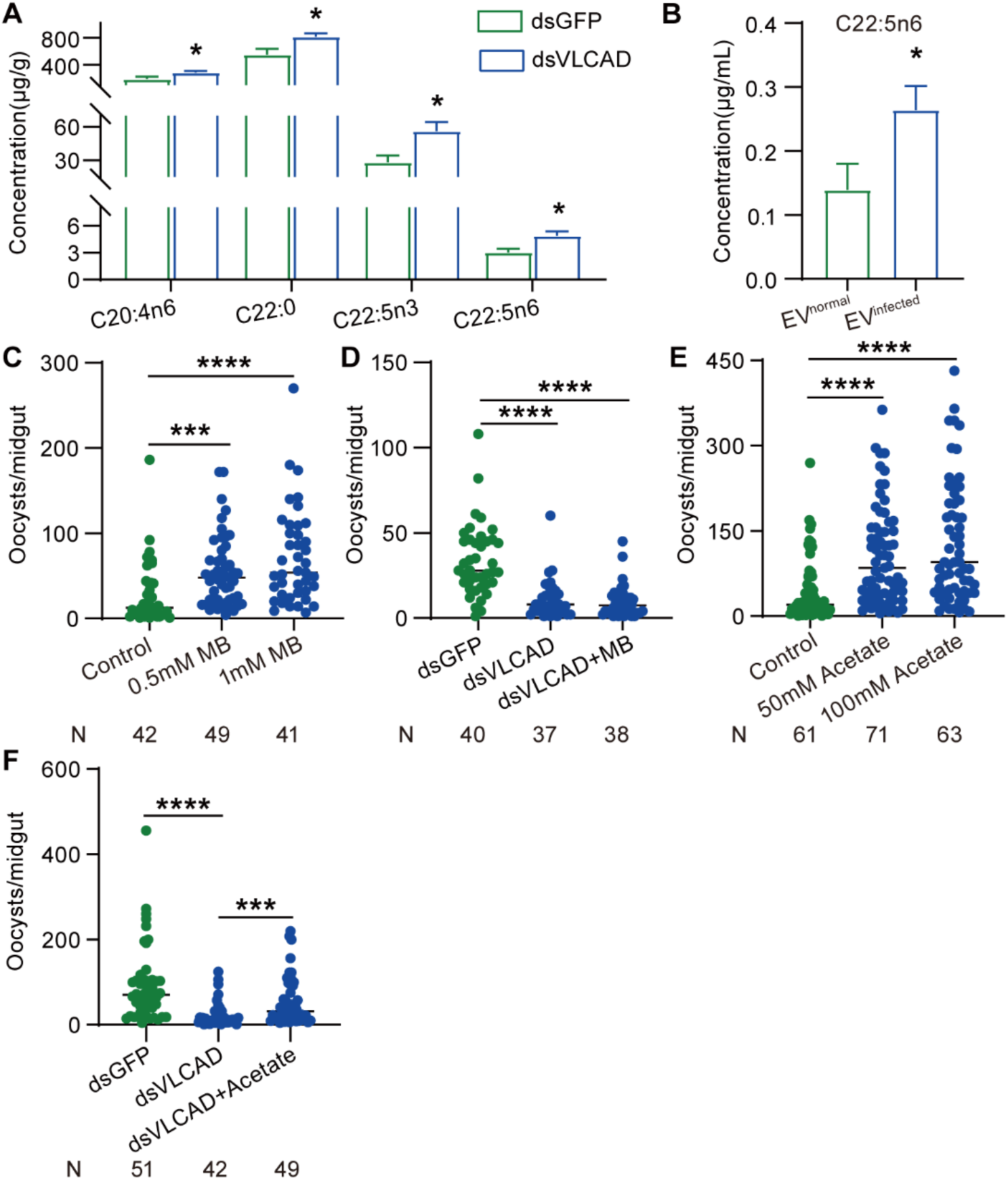
Fatty Acid β-Oxidation via VLCAD Promotes *Plasmodium* Infection. **A**, Fatty acids profile of mosquitoes treated with dsGFP and dsVLCAD 48 hrs post normal blood. Data are presented as the mean ± SEM. **B**, Fatty acids profile of mosquito midguts treated with EV^normal^ and EV^infected^. Data are presented as the mean ± SEM. **C**, Oocyst numbers of mosquitoes orally supplemented with MB. **D**, Oocyst numbers of mosquitoes treated with dsGFP, dsVLCAD and dsVLCAD orally supplemented with 0.5 mM MB. **E**, Oocyst numbers of mosquitoes orally supplemented with acetate. **F**, Oocyst numbers of mosquitoes treated with dsGFP, dsVLCAD and dsVLCAD orally supplemented with 50 mM acetate. For (C) to (F), data were pooled from two independent experiments, and each dot represents an individual mosquito. N, number of mosquitoes analyzed. The horizontal black lines indicate the median number of oocysts. Significance was determined by two-sided Student’s *t* test in (A) and (B) and ANOVA with Dunn’s test in (C), (D), (E) and (F). \**p* < 0.05, \*\*\**p* < 0.001, \*\*\*\**p* < 0.0001.

To determine whether the VLCFAs accumulation promotes *Plasmodium* infection, we supplemented mosquitoes with methyl behenate (C22:0, MB) via sugar feeding and quantified oocyst numbers. MB significantly increased oocyst burdens (Fig. 3C), but this effect was abolished in *VLCAD* knockdown mosquitoes (Fig. 3D), demonstrating that VLCAD activity is required for VLCFA–mediated enhancement of parasite infection.

Following VLCAD-mediated oxidation, VLCFAs are degraded into shorter acyl-CoA units and ultimately converted to acetyl-CoA through successive rounds of fatty acid β-oxidation^13^. To determine whether acetyl-CoA mediates the infection-promoting effect, we orally administrated acetate, a direct precursor of acetyl-CoA^14^, to mosquitoes for 24 hrs before infection. Acetate treatment significantly increased oocyst numbers (Fig. 3E). Moreover, acetate administration restoring oocyst numbers of *VLCAD* knockdown mosquitoes to control levels (Fig. 3F). Collectively, these results indicate that EV^infected^ enhances *Plasmodium* infection by stimulating VLCAD-dependent catabolism of VLCFAs to acetyl-CoA in the mosquito midgut.

### Acetyl-CoA Fuels Parasite Infection via Fatty Acid Biosynthesis

Acetyl-CoA is a central metabolic intermediate that feeds into diverse pathways, including protein acetylation, the tricarboxylic acid (TCA) cycle, and fatty acid synthesis^15,16^. To examine how acetyl-CoA promotes *Plasmodium* infection, we first tested whether this effect was mediated through histone acetylation. If so, inhibition of acetylation would be expected to reduce infection, whereas enhanced acetylation would increase parasite load. To test this, we inhibited acetylation either by RNAi-mediated knockdown of a mosquito histone acetyltransferase (HAT) or by oral administration of anacardic acid (AA), a HAT inhibitor. Conversely, we enhanced acetylation by knocking down histone deacetylase (HDAC) or by treating mosquitoes with trichostatin A (TSA), a HDAC inhibitor. Unexpectedly, inhibition of histone acetylation via *HAT* knockdown or AA treatment significantly increased oocyst numbers (Fig. S5A to S5D), while enhanced acetylation by *HDAC* knockdown or TSA treatment reduced parasite infection (Fig. S5E to S5H). These findings indicate that acetyl-CoA promotes *Plasmodium* infection through a mechanism independent of mosquito histone acetylation.

We next asked whether acetyl-CoA promotes infection by fueling the TCA cycle. Mosquitoes were supplemented with pyruvate or succinate, two key TCA cycle intermediates, and then challenged with *P. berghei*. Surprisingly, both treatments significantly reduced oocyst numbers, excluding a role for TCA fueling (Fig. S5I and S5J).

Acetyl-CoA also serves as a precursor for *de novo* fatty acid synthesis through conversion to malonyl-CoA^17^. To examine whether this pathway contributes to parasite infection, we supplemented mosquitoes with malonate, a precursor of malonyl-CoA. Malonate administration significantly increased oocyst numbers in *An. stephensi* (Fig. 4A), and importantly, also enhanced infection in *VLCAD* knockdown mosquitoes (Fig. 4B), indicating that fatty acid synthesis promotes parasite development.

**Fig. 4.**
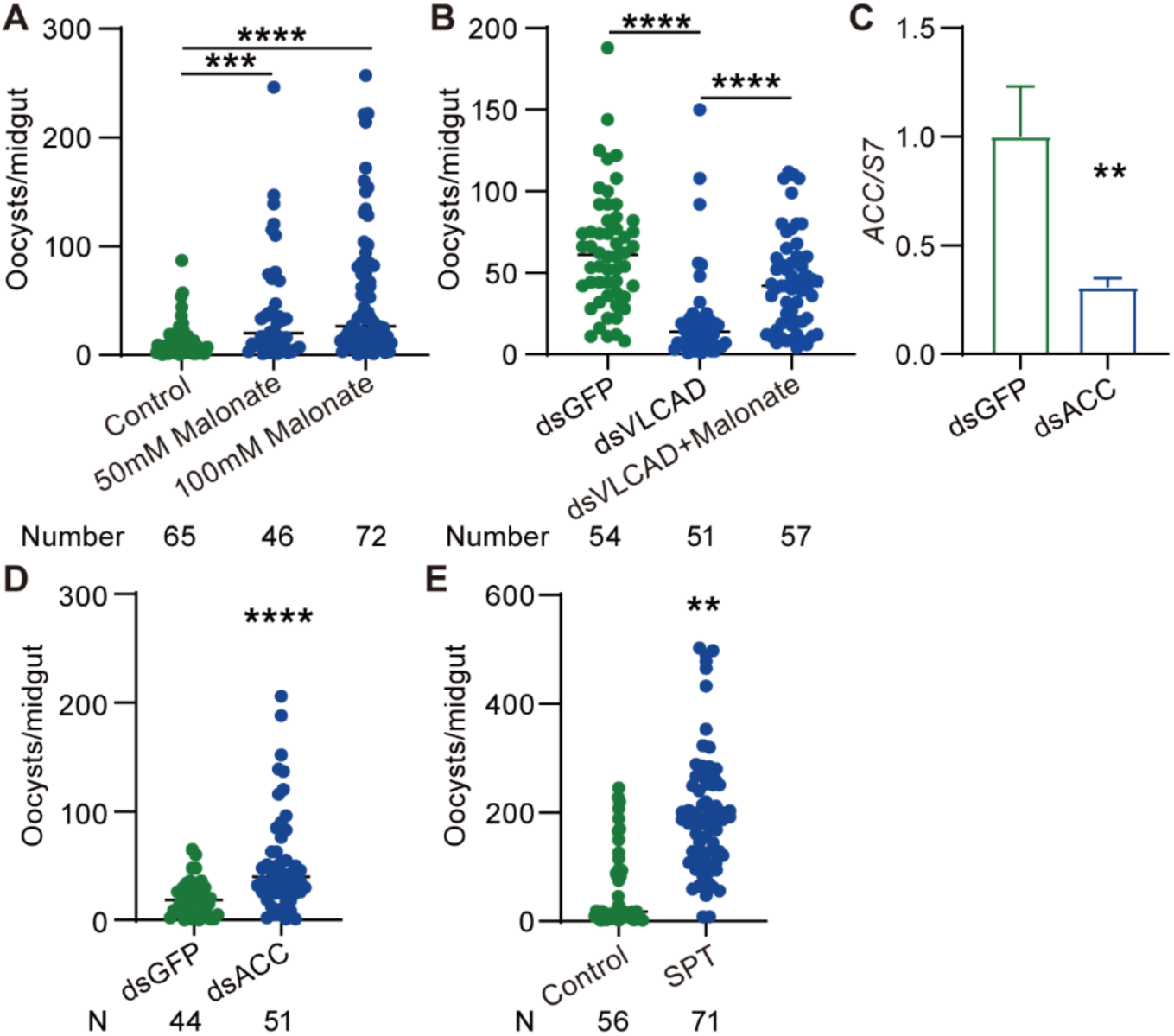
Malonyl-CoA promotes *Plasmodium* infection. **A**, Oocyst numbers in mosquitoes orally supplemented with malonate. **B**, Oocyst numbers of mosquitoes treated with dsGFP, dsVLCAD and dsVLCAD orally supplemented with 50 mM malonate. **C**, Knockdown efficiency of *ACC* in mosquitoes. The expression level of *ACC* was normalized to *An. stephensi S7*. Relative expression level of *ACC* in dsACC mosquitoes was normalized to that in dsGFP controls. Data are presented as the mean ± SEM (n=10 in dsGFP, n=10 in dsACC). **D**, Oocyst numbers of dsACC and dsGFP-treated mosquitoes. **E**, Oocyst numbers of mosquitoes treated with 5 mM SPT. For (A), (B), (D) and (E), data were pooled from two independent experiments, and each dot represents an individual mosquito. N, number of mosquitoes analyzed. The horizontal black lines indicate the median number of oocysts. Significance was determined by ANOVA with Dunn’s test in (A) and (B), two-sided Student’s *t* test in (C) and Mann-Whitney test in (D) and (E). \*\**p* < 0.01, \*\*\**p* < 0.001, \*\*\*\**p* < 0.0001.

In addition to usurping mosquito’s malonyl-CoA, *Plasmodium* also has the capability to synthesize its own malonyl-CoA^18^. To investigate whether *Plasmodium* depends on mosquito-derived malonyl-CoA for fatty acid synthesis, we disrupted malonyl-CoA production by knocking down *acetyl-CoA carboxylase* (*AC*C), which catalyzes the conversion of acetyl-CoA to malonyl-CoA. Knockdown of *ACC* significantly increased oocyst numbers (Fig. 4C and 4D). Similarly, treatment with spirotetramat (SPT), a potent ACC inhibitor^19^, enhanced parasite infection (Fig. 4E). These results suggest that inhibition of mosquito malonyl-CoA synthesis causes acetyl-CoA accumulation, which is scavenged by the parasite for its own malonyl-CoA production. Together, these findings demonstrate that acetyl-CoA enhances *Plasmodium* infection not through acetylation or the TCA cycle, but by fueling fatty acid synthesis. Both acetyl-CoA and malonyl-CoA can be usurped by parasites to sustain their metabolic needs in the mosquito midgut.

### *Plasmodium* Harnesses Mosquito β-Oxidation to Fuel Its Own Lipid Synthesis

To verify that acetyl-CoA produced from MB β-oxidation contributes to parasite lipid synthesis, we performed stable isotope tracing experiments. Mosquitoes were fed [1,2,3,4-¹³C₄]-behenic acid (BA) for 24 hrs prior to *P. berghei* infection. Ten days post infection, individual oocysts were isolated, and non-targeted metabolomics by particle-enhanced laser desorption/ionization mass spectrometry (PELDI-MS) was performed (Fig. 5A). Labeled carbons from [¹³C_4_]-BA were incorporated into multiple fatty acids, including C20:5, C20:0, C18:3, C18:2, C18:1, C18:0, C16:1, C15:0, C10:0 compared to controls (Fig. 5B). Together with our finding that MB supplementation failed to enhance oocyst numbers in *VLCAD* knockdown mosquitoes (Fig. 3D), these results demonstrate that *Plasmodium* exploits the mosquito fatty acid β-oxidation pathway to generate acetyl-CoA, which is then redirected into parasite fatty acid biosynthesis to sustain oocyst development.

**Fig. 5.**
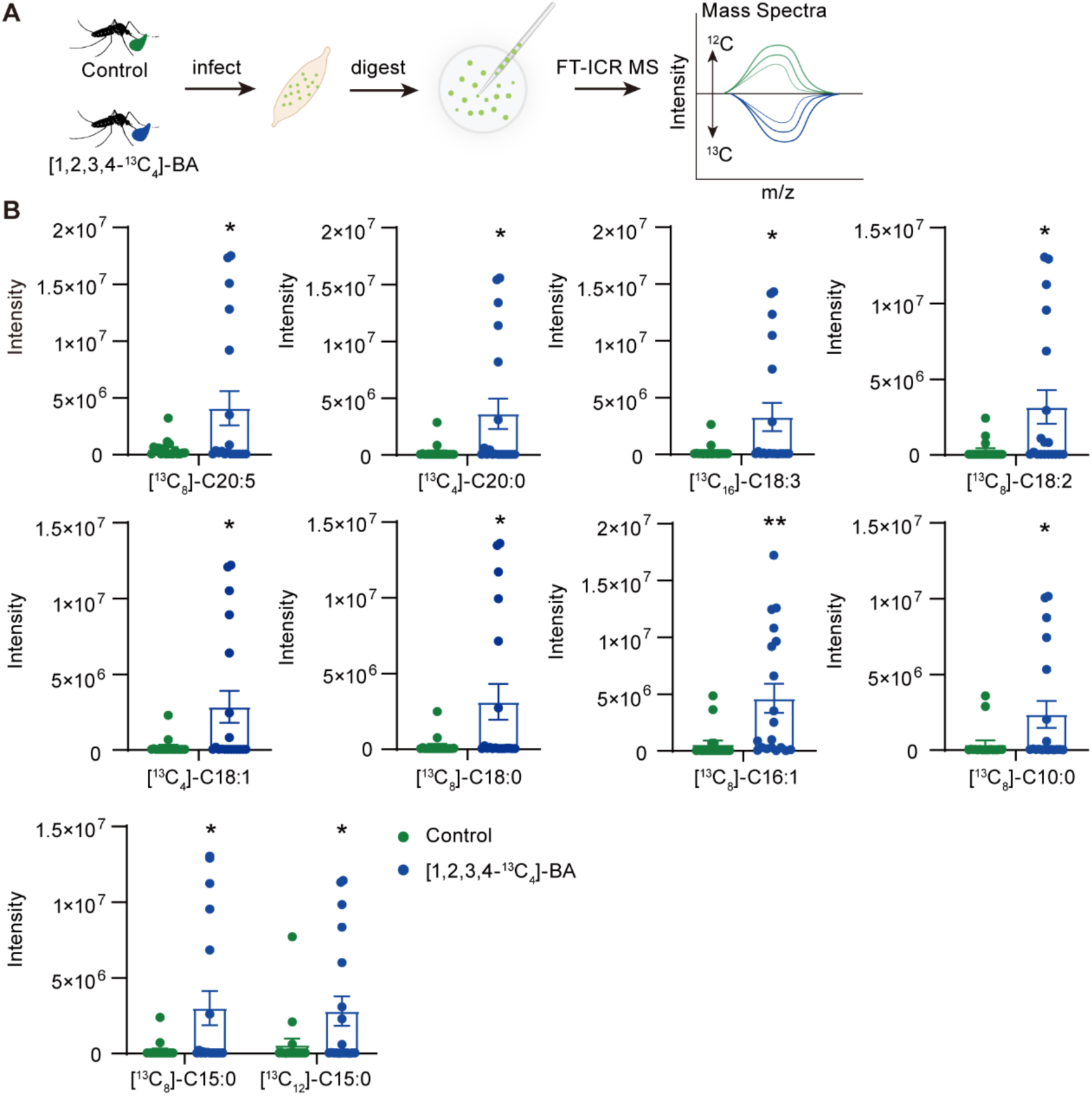
Plasmodium scavenges acetyl-CoA from mosquito to synthisize fatty acids. **A,** The workflow of isotope tracing analysis of *Plasmodium* oocyst lipids. **B**, Intensity of^13^C-labelled fatty acids of *P. berghei* oocysts. Data are presented as the mean ± SEM (n=18 in vehicle treated, n=19 in [1,2,3,4-^13^C_4_]-BA treated). Significance was determined two-sided Student’s *t* test. \**p* < 0.05, \*\**p* < 0.01.

### Ceramides enriched in host EVs promotes oocyst development

The elevated levels of VLCAD and C22:5n6 fatty acids in mosquito midguts following EV^infected^ treatment (Fig. 2C and 3B) prompted us to examine EVs lipid composition. Lipidomic profiling of EV^normal^ and EV^infected^ derived from mice by liquid chromatography–mass spectrometry (LC–MS) identified 70 lipid species, with multiple ceramides markedly enriched in EV^infected^. These included hexosylceramide, (Hex1Cer) (d18:1/16:0, d18:1/18:0, d18:0/24:0, d18:1/20:0, d18:1/24:0, d18:1/22:0), dihexosylceramides, (Hex2Cer) (d18:1/16:0), dihexosylceramides, (DHCer) (d18:0/16:0), and Cer (d18:1/22:1) (Fig. 6A and 6B). This finding is consistent with prior observations that EVs from *P. falciparum*–infected red blood cells are ceramides-enriched^20^. To examine whether these ceramides are transferred to mosquitoes, we measured midgut ceramide levels 24 hrs after EVs exposure using anti-ceramide immunostaining. EV^infected^ –treated mosquitoes displayed stronger signals than EV^normal^ controls (Fig. 6C). Similarly, feeding on *P. berghei*–infected mice resulted in pronounced ceramides accumulation in the midgut of mosquitoes, compared to feeding on non-infected mice (Fig. 6D). We next examined whether exogenous ceramide could promote *Plasmodium* infection. As expected, supplementation of Cer (d18:1/22:0) in blood meals increased oocyst burden (Fig. 6E and 6F). In parallel, MB feeding also elevated parasite loads (Fig. 6E and 6G).

**Fig. 6.**
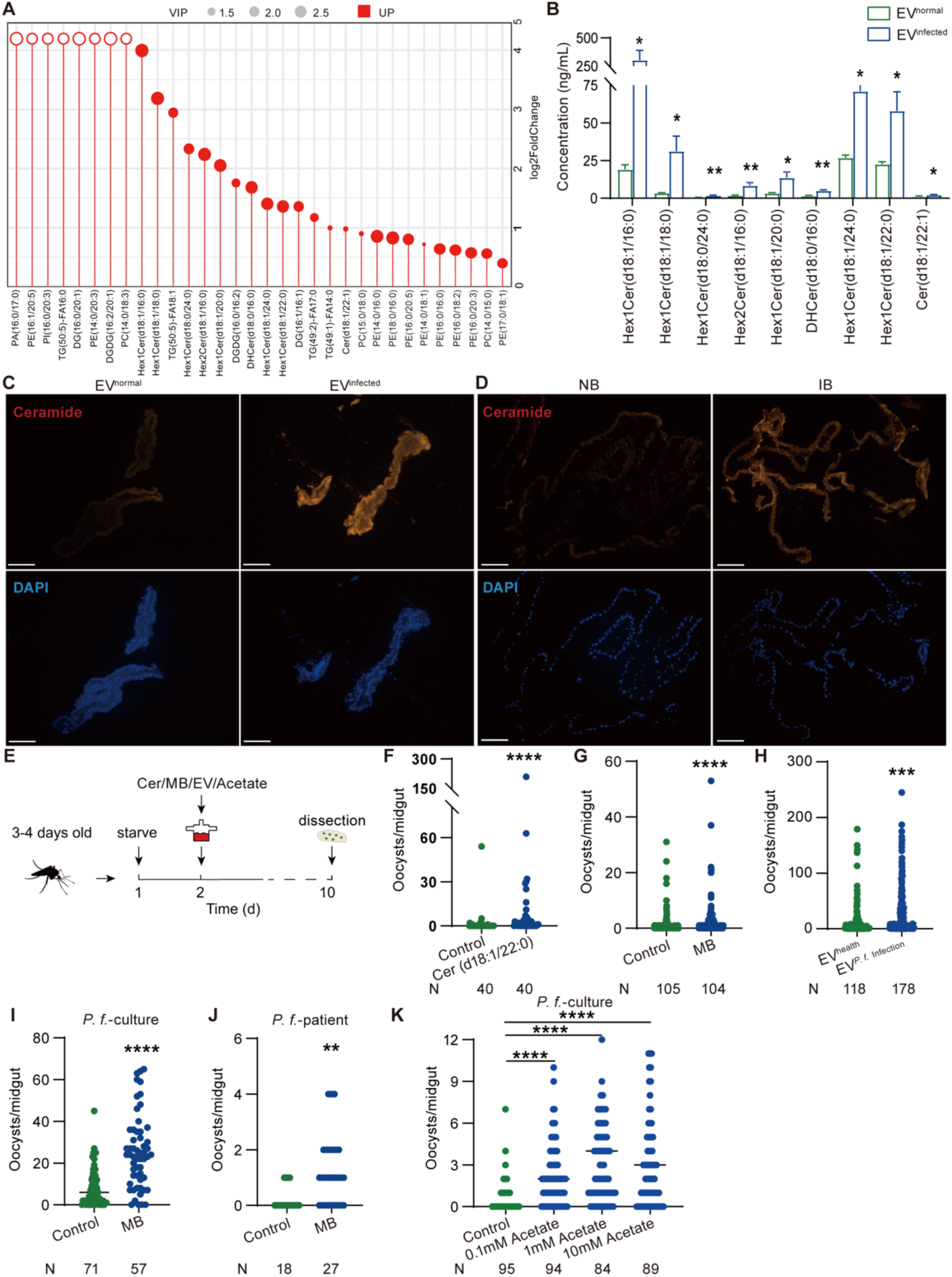
Ceramides accumulated in EV^infected^ promote *Plasmodium* transmission to mosquitoes. **A**, The enriched lipids in EV^infected^ compared to EV^normal^. Hollow circle represents log2foldchang > 5. **B**, Quantification of ceramides in EV^normal^ and EV^infected^. **C and D,** Immunostaining of ceramides (red) in the midgut of mosquitoes treated with EV^normal^ and EV^infected^ prior to infection (**C**) and 24 hrs post NB and IB (**D**). Nucleis were stained by DAPI (blue). Scale bar, 100 μm. **E**, Workflow of mosquito treatment using SMFA. **F and G**, Oocyst numbers of mosquitoes supplemented with 10 μM Cer (d18:1/22:0) (**F**) and 10 μM MB (**G**) through SMFA. **H**, Oocyst numbers of mosquitoes orally supplemented with EV^health^ derived from health human sera and EV*^P.f^*^infected^ derived from *P. falciparum* patient sera. **I and J**, Oocysts numbers of *P. falciparum* derived from culture (**I**) and from malaria patient (**J**) in mosquito treated with 10 μM MB. **K,** Oocyst numbers of *P. falciparum* of mosquito treated with acetate. For (F) to (K), data were pooled from at least two independent experiments and each dot represents an individual mosquito. N, number of mosquitoes analyzed. The horizontal black lines indicate the median number of oocysts. Significance was determined by two-sided Student’s *t* test in (B), Mann-Whitney test in (F) to (J) and ANOVA with Dunn’s test in (K). \**p* < 0.05, \*\**p* < 0.01, \*\*\**p* < 0.001, \*\*\*\**p* < 0.0001.

To assess whether this phenomenon extends across *Plasmodium* species, we administered EVs derived from *P. falciparum*–infected patient blood (EV*^P.f^*^infected^) to mosquitoes and challenged them with *P. berghei*. EV*^P.f^*^infected^ significantly boosted oocyst numbers compared to EV^health^ controls (Fig. 6H). Moreover, supplementation with MB increased the oocyst numbers of *P. falciparum* derived from culture (Fig. 6I) and notably, also enhanced infection of *P. falciparum* derived from malaria patient in mosquitoes (Fig. 6J). As expected, acetate supplementation similarly increased *P. falciparum* loads (Fig. 6K). Altogether, these results indicate that ceramides-enriched in EV^infected^ from the mammalian host promote parasite transmission across species via mosquito fatty acid β-oxidation pathway.

## Discussion

EVs secreted by host cells play central roles in *Plasmodium* invasion, pathogenesis, and drug resistance within mammalian hosts^8–10^. During blood feeding, these EVs are co-ingested with parasites into the mosquito midgut, but whether they function cross species to influence infection remains unclear. Here we show that EVs derived from *Plasmodium*-infected mouse and human serum enhance *Plasmodium* infection in mosquitoes by remodeling mosquito fatty acid catabolism. We further identify ceramides, enriched in EVs from infected hosts, as the key activators of this metabolic reprogramming in mosquitoes.

The role of EVs in mediating pathogen-pathogen and pathogen–host interactions within species has been well established^21^. Increasing evidence also indicates that EVs facilitate cross-species communication during vector-borne pathogen transmission, particularly from insect vectors to mammalian hosts. For example, EVs from mosquito saliva deliver subgenomic flaviviral RNA or sphingomyelins lipids to human cells, enhancing viral replication by suppressing interferon responses or ER-associated degradation of viral protein^22,23^. *Leishmania* parasites co-egest EVs with the parasite during sand fly biting, significantly increasing host infection rates^24^. By contrast, the contribution of mammalian host–derived EVs to pathogen transmission into insect vectors remains largely unexplored. Here, we demonstrate that EVs from *Plasmodium*-infected serum are specifically internalized by mosquito midgut epithelial cells, where they promote parasite oocyst development. Although EVs from uninfected hosts are similarly taken up, their physiological effects on mosquito biology remain unclear and warrant further study.

*Plasmodium* infection drives the packaging of parasite-derived molecules, including small RNAs, genomic DNA, and proteins, into EVs, which can manipulate host immune responses, vascular function, and cytoskeletal organization to favor parasite development^9,10,25,26^. Our work further demonstrates that *Plasmodium* remodels the lipidomic profile of host EVs, enriching them with ceramides.When ingested by mosquitoes, these EVs upregulate VLCAD, thereby boosting acetyl-CoA production, which is essential for oocyst development^27^. Given that overall lipid levels are reduced in the sera of *Plasmodium*-infected mice and humans^28,29^, and that blood-derived lipids are critical for the growth of both mosquitoes and parasites^30–32^, it is possible that *Plasmodium* manipulates host lipid metabolism by packaging lipids such as ceramides into EVs. This strategy likely compensates for lipid depletion in the infected host while simultaneously promoting parasite development in the mosquito vector. Beyond ceramides, the roles of other EVs-derived molecules in mediating *Plasmodium* transmission from host to vector remain poorly understood and warrant further investigation.

The oocyst stage of *Plasmodium* is highly metabolically demanding, characterized by repeated mitotic divisions that generate thousands of sporozoites^33–36^. However, oocyst metabolism remains poorly understood, largely due to challenges in culturing oocysts *in vitro* and isolating them from mosquito midguts. In this study, we established a metabolomic assay capable of analyzing as few as 10 oocysts directly isolated from mosquito midguts. Using stable isotope tracing, we show that long-chain fatty acids processed by mosquito β-oxidation are subsequently incorporated into oocyst lipids. Lipids are indispensable at this stage, serving as essential building blocks for membrane biogenesis^37^. As an obligate parasite, *Plasmodium* acquires fatty acids through both host scavenging and *de novo* fatty acid synthesis (FAS) during infection of mammalian hosts mammalian hosts^38–44^. For example, *P. falciparum* ribosomal protein P2 facilitates uptake of oleic and palmitic acids from host sera to support schizogony^45^, and radiolabeled fatty acids supplied in culture medium are efficiently incorporated into intraerythrocytic *P. falciparum* phospholipids and neutral lipids^42^. Parasites also scavenge fatty acids by hydrolyzing host lysophosphatidylcholine^46^. Genetic evidence further confirms that parasite FAS and fatty acid elongation (FAE) pathways are essential during liver stage development^39–44,47^. Our findings extend this model by showing that *Plasmodium* scavenges acetyl-CoA and malonyl-CoA from mosquitoes to fuel its own fatty acid synthesis. This aligns with the previous observations that *Plasmodium* spp. lacks a complete classical fatty acid β-oxidation pathway^48^. Consistently, mutations in parasite FAS or FAE genes, including *FabB/F*, *FabI*, and 3-hydroxyacyl-CoA dehydratase, impair oocyst maturation and sporozoite formation^43,44,47^. Recently single-cell sequencing analyses further confirm the active lipid biosynthesis during oocyst developments^49^. Together, these findings indicate that oocysts exploit mosquito fatty acid oxidation to generate acetyl-CoA, which serves as a substrate for fatty acid synthesis and elongation.

In summary, *Plasmodium* has evolved intricate strategies to adapt to the contrasting environments of the mammalian host and mosquito vector. By remodeling host EVs lipid composition, the parasite activates mosquito fatty acid β-oxidation to supply acetyl-CoA, which is essential for oocyst development. Targeting acetyl-CoA production in mosquitoes may thus provide a promising strategy to block malaria transmission.

## Methods

### Ethics statement

Collection of blood samples from healthy donors and malaria patients was approved by the Institutional Review Board of the Shandong Institute of Parasitic Diseases, China (SXB202410). All samples were obtained for routine diagnostic tests, with no additional burden to participants. Written informed consent was obtained from all healthy donors and malaria patients for the collection and research use of biological materials. All animal experiments were conducted in accordance with the guidelines for animal care and use of Fudan University and were approved by the Animal Care and Use Committee, Fudan University, China.

### Mosquito maintenance and *Plasmodium* infection

*An. stephensi* adults were maintained at 28°C, 80 ± 5% relative humidity, under a 12 hrs light/dark cycle, and provided 10% sucrose adlibitum. BALB/c mice were used to maintain mosquito colonies.

For *P. berghei* infections, starved female *An. stephensi* were allowed to feed on BALB/c mice with *P. berghei* (ANKA) parasitemia of 3–4%. Control and treatment groups were fed on the same mouse in parallel. After feeding, mosquitoes were maintained at 21°C and supplied daily with 10% sucrose. Non-engorged females were removed at 24 hrs post–blood meal. To quantify ookinetes, midguts were dissected 24 hrs post infection, blood boluses removed, and preparations mounted under a coverslip. Ookinetes were counted at 200× magnification using a Nikon Eclipse Ni-U microscope. For oocyst quantification, midguts were dissected 8 days post infection and examined at 100 × magnification. For sporozoite quantification, salivary glands were dissected 21 days post infection, homogenized in 0.9% NaCl, and sporozoites were counted using a hemocytometer at 400 × magnification.

For *P. falciparum* infection, *P. falciparum* strain NF54 was kindly provided by the Marcelo Jacobs-Lorena laboratory at Johns Hopkins University and maintained under standard culture conditions^50,51^. Briefly, parasites were revived using a prewarmed revival solution (16% D-Sorbitol, 0.9% NaCl) maintained at 37°C, with a hematocrit of 2%. Parasites were maintained in RPMI 1640 medium (Gibco) supplemented with 10 mg/L hypoxanthine, 20 mM HEPES, 10% human serum, and L-glutamine. The parasite ring stage was diluted to 0.5%, maintaining a hematocrit of 4%. After continuous culture for 15 days, mature gametocytes were precipitated by centrifugation (1,000 × g, 5 min). Infections were performed using the Standard Membrane Feeding Assay (SMFA). Gametocytes were added into RBCs at a final concentration of 0.15%, and then supplemented with an equal volume of human serum to reconstitute whole blood, maintained at 37°C. For *P. falciparum* infection using patient blood, blood was collected from *P. falciparum*-malaria patients into 10 mL heparinized tubes. The blood was then fed to starved mosquitoes at 37°C for 30 min using SMFA. Mosquitoes were then cold-anesthetized. Unfed mosquitoes were removed. Infected mosquitoes were maintained at 27°C for 8 days. Then midguts were dissected and stained with 0.2% mercurochrome. Oocyst were counted microscopically.

To assess blood feeding capacity and reproduction, mosquitoes were weighed immediately after blood feeding. At 48 hrs post feeding, ovaries were dissected, and eggs were counted microscopically.

### EVs isolation and administration

Sera-derived EVs were isolated from BALB/c mice or humans as described previously^52,53^. Briefly, blood was collected from *P. berghei*-infected (parasitemia 3-4%) and uninfected mice or from *P. falciparum*-infected patients and health donors. Samples was incubated at room temperature for 30 min, then centrifuged at 4°C under the following conditions: 300 × g for 10 min, 2,000 × g for 10 min, and 10,000 × g for 30 min. Serum fractions were pooled after each step, filtered through a 0.22-μm filter, and centrifuged at 150,000 × g for 3 hrs. Pellets were resuspended in 1 × phosphate-buffered saline (PBS) and subjected to ultracentrifugation at 150,000 × g for an additional 3 hrs to remove contaminants. Finally, EVs pellets were resuspended in 1 × PBS and stored at −80°C until use. EV morphology was assessed by TEM, and size distribution was analyzed by NTA. For mosquito feeding assays, EVs pellets were dissolved in 1.5% sucrose and orally administered to starved mosquitoes for 24 hrs.

### EVs internalization assay

EVs pellets were suspended in 250 μL Diluent C and labelled with 2 μL PKH67 dye (Sigma) according to the manufacturer’s instructions. Staining was quenched with 1% BSA for 1 min to bind excess dye. The mixture was filtered through a 100 kDa MWCO Amicon® Ultra column by centrifugation at 4,000 × g for 10 min at room temperature, and the process was repeated several times until the entire mixture had passed through. EVs were collected by ultracentrifugation at 150,000 × g for 3 hrs at 4°C and resuspended in PBS. For *in vitro* analysis, MSQ43 cells were grown in Schneider’s medium (Gibco) supplemented with 10% heat-inactivated fetal bovine serum (FBS, Gibco), 100 IU/mL penicillin and 100 μg/mL streptomycin (Thermo Fisher) at 28°C. PKH67 labelled-EVs were added to 1 × 10^6^ MSQ43 cells. After 24 hrs of incubation, cells were washed three times with PBS and fixed in 4% paraformaldehyde for 30 min at room temperature. Nuclei were stained with DAPI, and EVs internalization was visualized using a Nikon ECLIPSE IVi microscope. PKH67 dye alone (no EV) was used as a negative control. For *in vivo* analysis, PKH67-labelled EVs were suspended in 1.5% sucrose and orally administered to mosquitoes starved for 24 hrs. The midguts, malpighian tubules, and ovaries were dissected and washed with PBS for 24 hrs post EVs administration. Nuclei were stained with DAPI, and EVs internalization was assessed using a Nikon ECLIPSE IVi microscope. Mosquitoes fed with PKH67 dye alone (no EV) served as negative controls.

### Proteome Profiling by Nano-LC–MS/MS

A total of 200 EVs-treated mosquito midguts were dissected on ice and washed twice with cold PBS. Tissues were lysed in TCEP buffer (2% sodium deoxycholate, 40 mM 2-chloroacetamide, 100 mM Tris-HCl, 10 mM tris(2-carboxyethyl) phosphine, 1 mM PMSF, and 1 mM protease/phosphatase inhibitor cocktail, pH 8.5) at 99°C for 30 min. After cooling to room temperature, proteins were digested with trypsin (Promega) at 37°C for 18 hrs. Digestion was terminated with 10% formic acid, followed by vortexing for 3 min and centrifugation at 12,000 × g for 5 min. The supernatant was transferred to a new 1.5 mL tube containing extraction buffer (0.1% formic acid in 50% acetonitrile), vortexed for 3 min, and centrifuged again at 12,000 × g for 5 min. The final supernatant was collected and analyzed by nano-LC–MS/MS.

Peptide samples were analyzed on a Q Exactive HF-X Hybrid Quadrupole-Orbitrap Mass Spectrometer (Thermo Fisher Scientific) coupled to an EASY-nLC 1200 system (Thermo Fisher Scientific). Dried peptides were dissolved in Solvent A (0.1% formic acid in water) and loaded onto a 2-cm self-packed trap column (100 μm inner diameter, 3 μm ReproSil-Pur C18-AQ beads), followed by separation on a 30-cm analytical column (150 μm inner diameter, 1.9 μm ReproSil-Pur C18-AQ beads) over a 150-min linear gradient from 4% to 100% Solvent B (0.1% formic acid in 80% acetonitrile) at a constant flow rate of 600 nL/min. Eluted peptides were ionized at 2 kV and analyzed in data-dependent acquisition mode. For MS1 full scans, ions with m/z 300–1,400 were acquired in the Orbitrap at a resolution of 120,000, with an automatic gain control (AGC) target of 3 × 10^6^ and a maximum injection time of 80 ms. The top 60 precursor ions were selected for higher-energy collisional dissociation (HCD) with normalized collision energy of 27%, and fragment ions were detected in the Orbitrap at a resolution of 7,500, with an AGC target of 5 × 10^4^ and maximum injection time of 20 ms. Dynamic exclusion of precursor ions was set to 25 s. Data were processed using Proteome Discoverer 2.3 (Thermo Fisher Scientific) for qualitative and quantitative analysis.

### RNA interference

Double-stranded RNAs (dsRNAs) targeting: *GAK3*, *VLCAD*, *ERGIC3*, *VPS13D*, *ACC*, *HAT*, *HDAC*, eGFP (BD Biosciences) were generated by PCR amplification of cDNA fragments using gene-specific primers (Table 1). The dsRNA was synthesed *in vitro* using the MEGAscript RNA kit (Thermo Fisher Scientific). Female *An. stephensi* mosquitoes (3–4 days old) were injected intrathoracically with 69 nL dsRNA (3–4 μg/μL) using a Nanoject II microinjector (Drummond). An equivalent amount of eGFP dsRNA was used as a control. Gene silencing efficiency was evaluated two days post-injection, and mosquitoes were challenged with infectious blood meals four days after microinjection.

### RNA extraction and quantitative PCR analysis

Total RNA was extracted from individual mosquitoes or from three pooled midguts using TRI reagent (Sigma) according to the manufacturer’s instructions. First-strand cDNA was synthesized from total RNA using the Hifair® III 1st Strand cDNA Synthesis SuperMix for qPCR (Yeasen). Gene expression levels were quantified on a LightCycler 96 Real-Time PCR System (Bio-rad) using SYBR Green qPCR Master Mix (Yeasen), and data were analyzed with the LightCycler 96 software. The ribosomal protein gene *S7* was used as an internal reference^54^, and relative expression was calculated using the 2^-ΔΔCt^ method^55^. Primer sequences are listed in Table 1.

### Western blot analysis

Ten midguts were pooled per sample and lysed in RIPA buffer containing complete protease inhibitor cocktail (Beyotime). Immunoblotting was performed by standard procedures using anti-VLCAD polyclonal antibody (1:2000, Abcam) and anti-Actin polyclonal antibody (1:2000, Abbkine), followed by HRP-conjugated secondary antibody (1:5000, Abbkine).

### Fatty Acid Profiling by GC–MS

To profile fatty acids, 50 mosquitoes treated with dsGFP and dsVLCAD 48 hrs post-blood meal, or 200 midguts from mosquitoes supplemented with EV^normal^ and EV^infected^, were pooled for one biological replicate. Six to eight replicates were analyzed by GC– MS. Samples were snap-frozen in liquid nitrogen, homogenized in 1 mL chloroform: methanol (2:1, v/v), and extracted ultrasonically for 30 min. Homogenates were centrifuged at 12,000 rpm for 5 min at 4°C, and supernatants were transferred to 15 mL tubes. Each sample was mixed with 2 mL of 1% sulfuric acid in methanol, vortexed for 1 min, and subjected to esterification. Fatty acid methyl esters were extracted with 1 mL n-hexane, vortexed, and allowed to stand for 5 min. Samples were then washed with 5 mL H₂O, centrifuged at 3,500 rpm for 10 min at 4°C, and supernatants were collected. Residual water was removed by adding 100 mg anhydrous sodium sulfate, followed by centrifugation at 12,000 rpm for 5 min at 4°C. A final 300 μL extract was transferred to a 2 mL tube, and 15 μL methyl salicylate (500 ppm) was added as an internal standard.

GC–MS analysis was performed on a Trace 1300 gas chromatograph (Thermo Fisher Scientific) equipped with a Thermo TG-FAME capillary column (50 m × 0.25 mm i.d., 0.20 μm film). Helium was used as the carrier gas at 0.63 mL/min. Samples were injected in split mode (8:1) with an injection volume of 1 μL at 250°C. The ion source and transfer line temperatures were set at 300°C and 280°C, respectively. The oven temperature program was as follows: initial 80°C (1 min), ramp to 160°C at 20°C/min (hold 1.5 min), ramp to 196°C at 3°C/min (hold 8.5 min), then ramp to 250°C at 20°C/min (hold 3 min). Mass spectra were acquired on an ISQ 7000 system (Thermo Fisher Scientific) using electron impact ionization (70 eV) in single ion monitoring (SIM) mode. Free fatty acids were identified and quantified using Thermo Scientific Chrome 7.2.10 software with integrated peak analysis and custom scripts.

### Metabolites and Inhibitors Supplementation in Mosquitoes

Metabolites supplementation: For supplementation via 10% sucrose: MB (Aladdin) was dissolved in 1% BSA by ultrasonication and dissolved in 10% sucrose. Sodium acetate, sodium pyruvate, sodium succinate and sodium malonate dibasic (Sigma-Aldrich) were dissolved individually in 10% sucrose. Then metabolites at indicated concentration were orally administered to starved mosquitoes (3–4 days old) for 24 hrs. Following an additional 24 hrs starvation, these mosquitoes were fed on *P. berghei*– infected mice. Midguts were dissected, and oocyst numbers were counted at 8 days post infection. For supplementation via blood: MB or sodium acetate were added into infected or normal blood and fed to starved mosquitoes using SMFA. Ceramide (Cer d18:1/22:0, Sigma) was dissolved in ethanol by ultrasonication on ice and added into infected or normal blood and fed to mosquitoes using SMFA.

Inhibitor supplementation: AA (Selleck), TSA (Selleck), and SPT (Aladdin) were dissolved in DMSO. Each compound was orally administered to starved mosquitoes in 10% sucrose for 24 hrs^19,56,57^. After an additional 24 hrs starvation, mosquitoes were fed on *P. berghei*–infected mice.

### Stable Isotope Tracing by FT-ICR Mass Spectrometry

Starved mosquitoes were treated with 2 mM [1,2,3,4-^13^C_4_]-BA (Sigma) for 24 hrs, followed by an additional 24 hrs of starvation prior to *P. berghei* infection. Midguts were dissected 10 days post infection and digested with 0.25% trypsin for 20–30 minutes as described^58^. Individual oocysts were collected using a glass capillary under fluorescence microscope. Ten oocysts were pooled into 2 μL 80% methanol, rapidly frozen in liquid nitrogen, and stored at –80°C for further metabolic analysis. For metabolic analysis, matrix particles were synthesized following established protocols^59^ and suspended at a final concentration of 1 mg/mL. Metabolite-containing samples were first loaded onto a microarray chip, followed by the addition of 0.3 μL of matrix suspension to each spot. The chip was then dried at room temperature for the following analysis.

Non-targeted metabolic analysis was performed using a Bruker Fourier-transform ion cyclotron resonance mass spectrometer (FT-ICR MS, SolariX 7.0T) in positive ion mode. The resulting mass spectra of the samples were collected. Raw spectral data were first centroided using Bruker Compass DataAnalysis software. Signals corresponding to even-numbered^13^C isotopologues (e.g., M+2) were included in the analysis. To identify and annotate potential downstream lipid metabolites derived from the^13^C-labeled precursor, accurate m/z values from FT-ICR-MS were matched against the Human Metabolome Database (HMDB) using a mass accuracy threshold of 10 ppm. Additionally, comparative analysis between labeled and unlabeled samples allowed for the quantification of label incorporation.

### EVs Lipid Extraction and LC–MS/MS Analysis

EVs from sera of eight normal or *Plasmodium*-infected mice were pooled per biological replicate, with six to eight replicates analyzed. Lipids were extracted using a modified MTBE method: samples were mixed with methanol containing internal standards, followed by MTBE addition, sonication, phase separation, and organic phase collection. Extracts were dried under nitrogen, reconstituted in isopropanol/acetonitrile, and clarified by centrifugation.

Lipidomic profiling was performed on an LC-30AD UHPLC system coupled to an AB 6500+ QTRAP mass spectrometer (AB SCIEX). Chromatographic separation was achieved using C18 and amino columns under optimized gradient conditions. Data were acquired in positive and negative ion modes with ESI and analyzed in MRM mode. Lipid identification and quantification were based on internal standards and peak integration.

### Immunostaining of ceramide

Mosquito midguts were dissected in cold PBS at 24 hrs post blood feeding or EVs adminstration, and fixed in 4% paraformaldehyde overnight at 4°C. Tissues were washed in PBS for 3 hrs, cryoprotected in a sucrose gradient, embedded in optimal cutting temperature (OCT) compound, and cryosectioned at 8 μm. Sections were permeabilized with 0.5% Triton X-100 in PBS (PBST) for 10 min and blocked with 10% FBS in PBST for 1 hr at room temperature. Samples were incubated with mouse anti-ceramide antibody (1:20; Sigma) overnight at 4°C, washed in PBS containing 0.05% Tween-20, and incubated with Alexa Fluor 555–conjugated anti-mouse secondary antibody (1:200; Thermo Fisher Scientific) for 2 hrs at room temperature. Nuclei were stained with DAPI, and samples were mounted in Fluoromount (Sigma). Images were acquired with a Leica SP8 confocal microscope.

## Statistics analysis

All analyses were performed in GraphPad Prism software (v.8). Statistical methods are specified in the Fig. legends. Comparisons between two groups were made using Student’s *t*-test (for gene expression and metabolite levels) or the Mann–Whitney test (for ookinetes, oocysts, and sporozoites). For multiple groups, one-way ANOVA followed by Dunnett’s (gene expression) or Dunn’s (parasite counts) test was applied.

## Acknowledgments

We acknowledgment our colleague, Jiong Ma, from the School of Information Science and Technology at Fudan University, for providing advice about isolating single *Plasmodium* oocyst.

## Author contributions

X.S., W.C., J.W. L.L., X.G., H.Z., H.Z., P.T., Q.X., K.Q., W.H., J.W. performed experiments and analyzed results. X.S., W.C., J.W. L.L., K.Q., W.H., J.W. conceived the experiments. X.S., W.C., J.W. L.L., K.Q., W.H., J.W. wrote and reviewed the manuscript. J.W., X.S. acquired Funding.

## Data and materials availability

All proteomic and lipidomics data have been deposited in Zenodo: 10.5281/zenodo.17168345. All the datasets generated and analyzed in the study are available in the main text and in the supplementary materials.

## Figure Legends

**Fig. S1.**
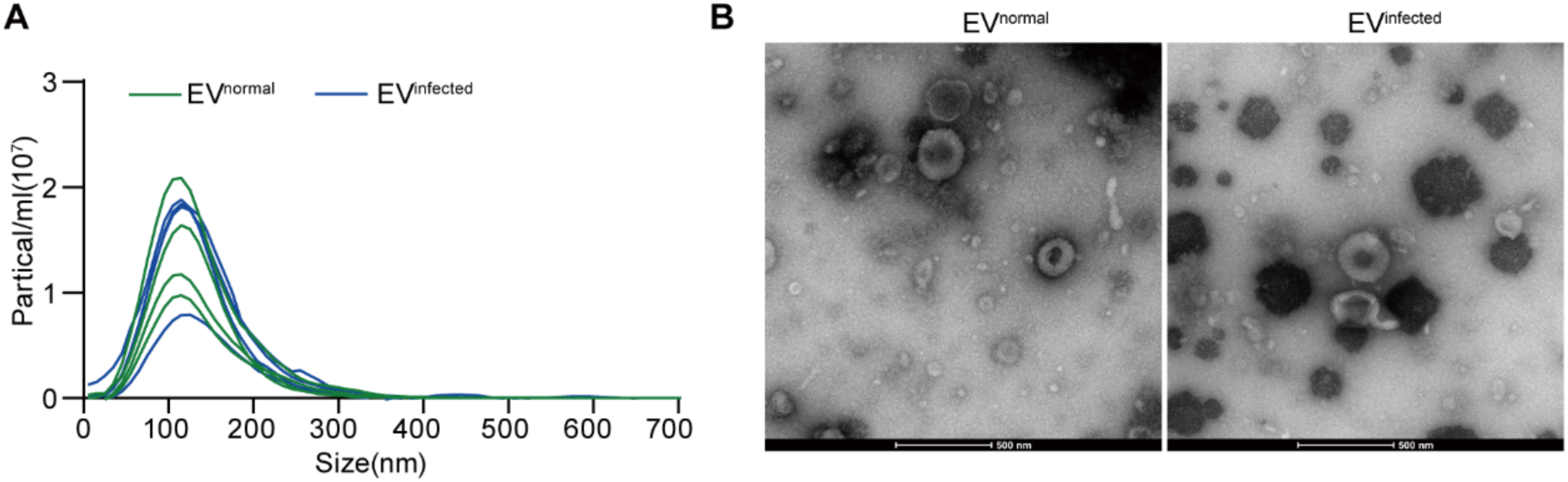
Concentration and morphology of EV^normal^ and EV^infected^. **A**, Nanoparticle tracking analysis (NTA) (size [nm] versus particles/mL) of EV^normal^ and EV^infected^. **B**, Transmission electron microscopy of EV^normal^ and EV^infected^. Scale bar: 500 nm.

**Fig. S2.**
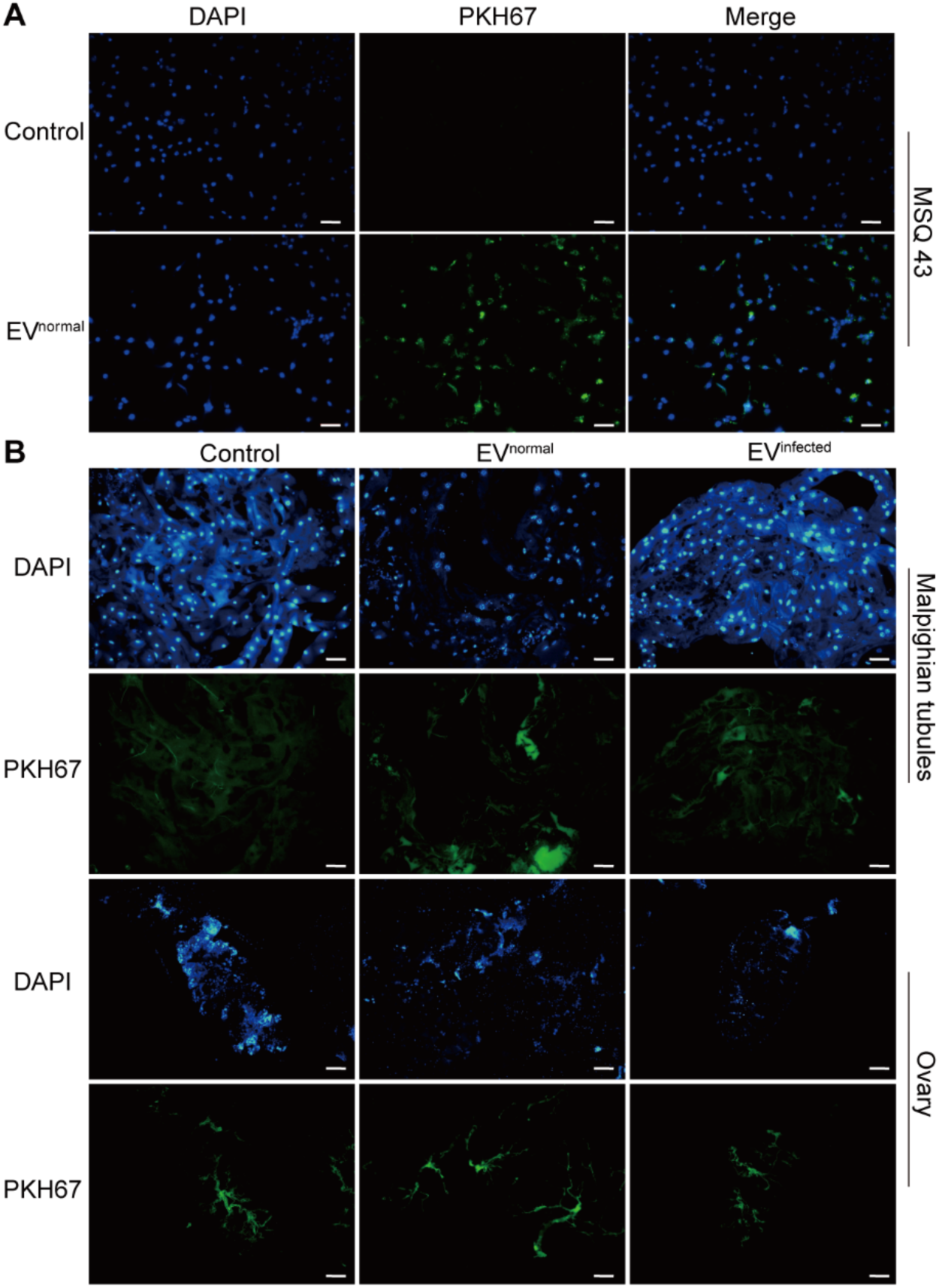
Internalization of EVs in mosquito cells and tissues. **A**, Internalization of PKH67-labelled EVs (green) in MSQ43 cells. **B**, Internalization of PKH67-labelled EVs (green) in malpighian tubules and ovaries. Nucleis were stained by DAPI (blue). Scale bar, 50 μm.

**Fig. S3.**
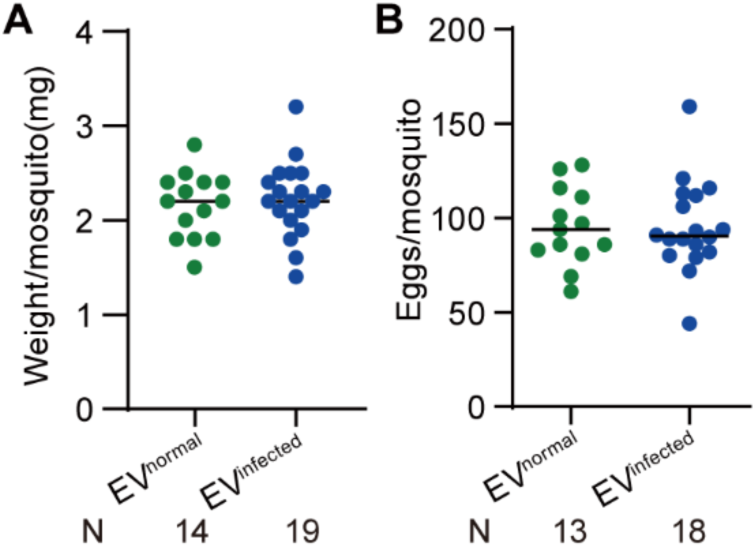
Influence of EVs treatment on mosquito blood feeding and reproduction. **A**, Weight of fully engorged mosquitoes treated with EV^normal^ and EV^infected^. **B**, Egg numbers of mosquitoes treated with EV^normal^ and EV^infected^. The horizontal black lines indicate the median, and each dot represents an individual mosquito in (A) and (B). Significance was determined by Mann-Whitney test.

**Fig. S4.**
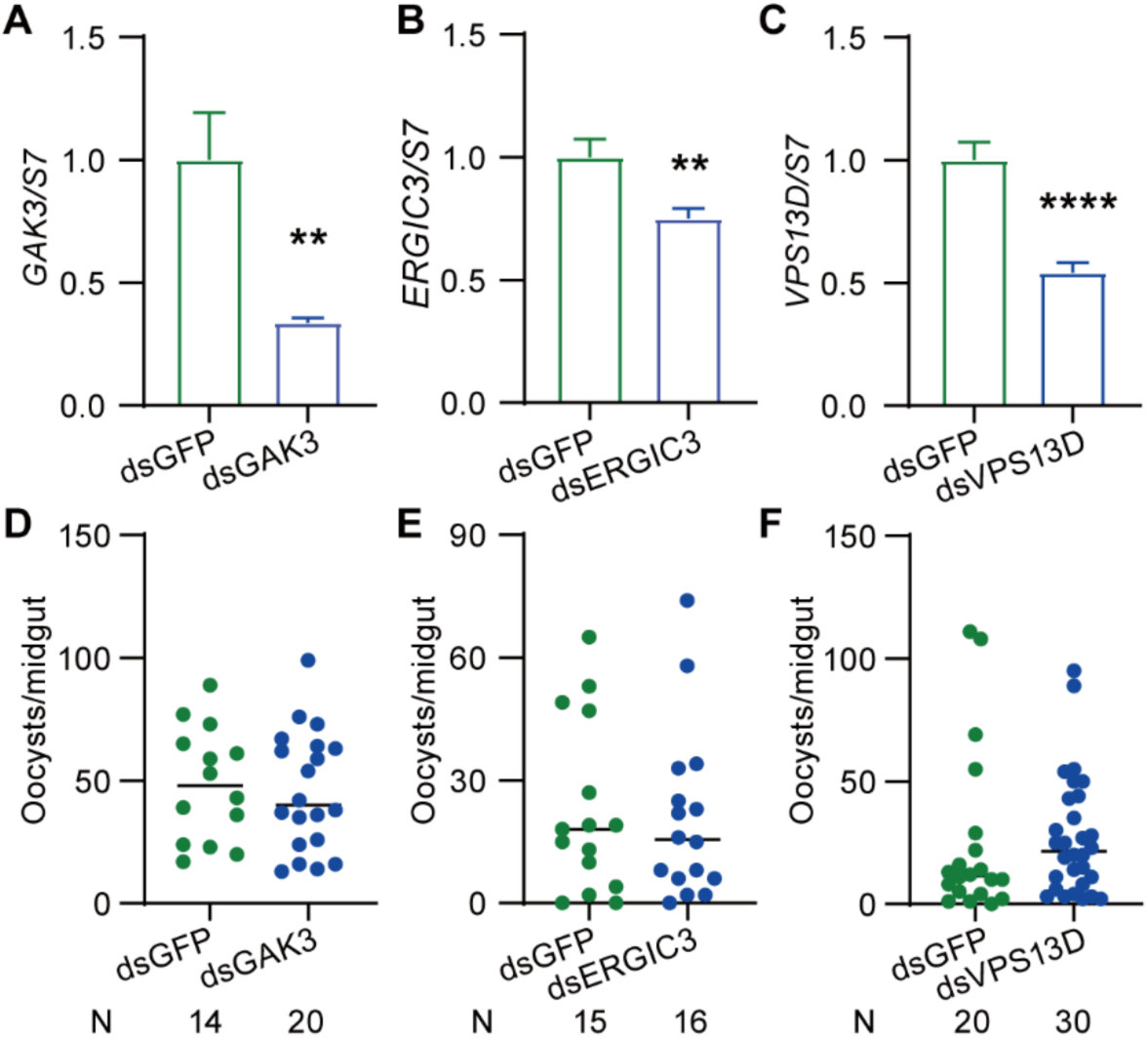
The influence of EV^infected^ –induced molecules on *Plasmodium* infection. **A to C**, Knockdown efficiency of *GAK* **(A)**, *ERGIC3* **(B)**, and *VPS13D* **(C)** in mosquitoes. The expression levels of the target genes were normalized to *An. stephensi S7*. Relative expression levels of these genes were normalized to those in dsGFP controls. Data are presented as the mean ± SEM (for A, n=10 in dsGFP, n=10 in dsGAK; for B, n=9 in dsGFP, n=10 in dsERGIC3; for C, n=10 in dsGFP, n=10 in dsVPS13D). **D** to **F**, Oocyst numbers of dsGAK **(D)**, dsERGIC3 **(E)**, dsVPS13D **(F)-treated** mosquitoes. For (D) to (F), each dot represents an individual mosquito. N, number of mosquitoes analyzed. The horizontal black lines indicate the median number of oocysts. Significance was determined by two-sided Student’s *t* test in (A) to (C) and Mann-Whitney test in (D) to (F). \*\**p* < 0.01, \*\*\*\**p* < 0.0001.

**Fig. S5.**
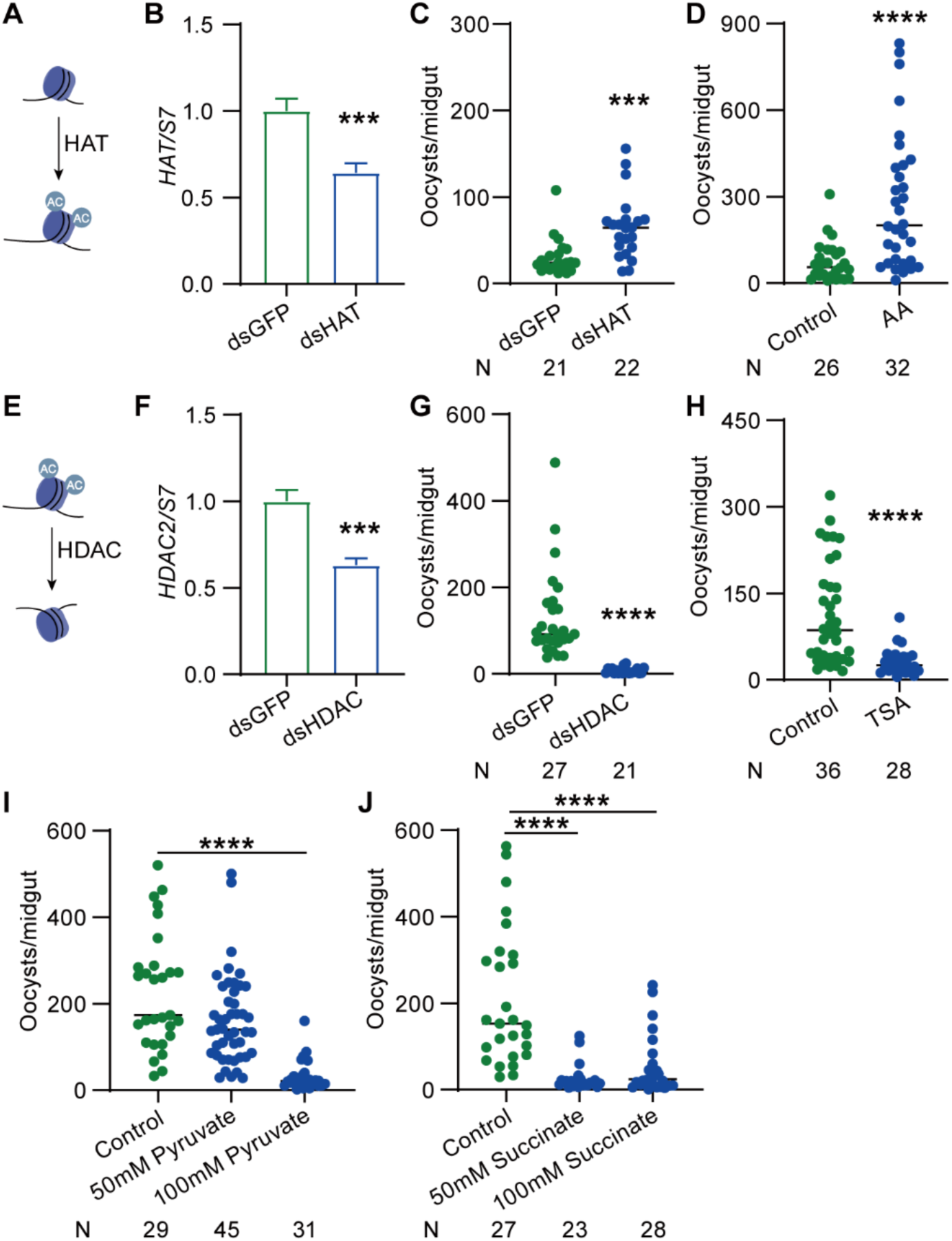
Acetyl-CoA facilitates *Plasmodium* infection in mosquitoes independent on acetylation and TCA cycle. **A and E**, Schematic diagram illustrating the functions of of HAT and HDAC functions**. B and F**, Knockdown efficiency of *HAT* (**B**) and *HDAC* (**F**) in mosquitoes. The expression levels of target genes was normalized to *An. stephensi S7*. Relative expression levels of target genes in dsRNA-treated mosquitoes were normalized to that in dsGFP controls. Data are presented as the mean ± SEM (for B, n=10 in dsGFP, n=10 in dsHAT; for F, n=10 in dsGFP, n=10 in dsHDAC). **C and G**, Oocyst numbers in the midgut of dsHAT (**C**) and dsHDAC (**G**)-treated mosquitoes. **D and H**, Oocyst numbers of mosquitoes orally supplemented with 10 μM AA (**D**) and 5 μM TSA (**H**). **I and J**, Oocyst numbers in mosquitoes orally supplemented with pyruvate (I) and succinate (J). For (C), (D), (G), (H), (I), and (J), each dot represents an individual mosquito. N, number of mosquitoes analyzed. The horizontal black lines indicate the median number of oocysts. Significance was determined by two-sided Student’s *t* test in (B) and (F), Mann-Whitney test in (C), (D), (G), and (H) and ANOVA with Dunn’s test in (I) and (J). \*\*\**p* < 0.001, \*\*\*\**p* < 0.0001.

